# *Coxiella burnetii*-containing vacuoles interact with host recycling-endosomal proteins Rab11a and Rab35 for vacuolar expansion

**DOI:** 10.1101/2024.03.01.582943

**Authors:** Brooke A. Hall, Kristen E. Senior, Nicolle T. Ocampo, Dhritiman Samanta

## Abstract

*Coxiella burnetii* is a gram-negative obligate intracellular bacterium and a zoonotic pathogen that causes human Q fever. Although acute Q fever manifests as atypical pneumonia, chronic infection may lead to life-threatening endocarditis. The lack of effective antibiotics and a licensed vaccine for *Coxiella* in the U.S. warrants further research into *Coxiella* pathogenesis. Within the host cells, *Coxiella* replicates in an acidic phagolysosome-like vacuole termed *Coxiella*-containing vacuole (CCV). Previously, we have shown that the CCV pH is critical for *Coxiella* survival and that the *Coxiella* Type 4B secretion system regulates CCV pH by inhibiting the host endosomal maturation pathway. However, the trafficking pattern of the ‘immature’ endosomes in *Coxiella*-infected cells remained unclear. Our recent CCV localization screen with host Rab proteins revealed that recycling endosome-associated proteins Rab11a and Rab35 localize to the CCV during infection, suggesting that CCV interacts with host recycling endosomes during maturation. Interestingly, only a subset of CCVs were Rab11a or Rab35-positive at any given time point. A quantitation of Rab11a/Rab35-positive CCVs at 3- and 6-days post-infection (dpi) revealed that at 3 dpi, ∼52% of CCVs were positive for Rab11a, whereas only ∼39% CCVs were positive for Rab35. This pattern reversed at 6-dpi, when only ∼22% of CCVs were positive for Rab11a and ∼64% of CCVs were positive for Rab35. These data suggest that the CCV preferentially interacts with Rab11a and Rab35-positive recycling endosomes depending on the stage of maturation. Furthermore, we observed a significant increase in Rab11a and Rab35 fluorescent intensity in *Coxiella-*infected cells compared to mock cells suggesting that *Coxiella* increases the recycling endosome content in infected cells. Finally, a siRNA-mediated knockdown of both Rab11a and Rab35 resulted in significantly smaller CCVs, suggesting that recycling endosomal Rab proteins are essential for CCV expansion. Overall, our data, for the first time, show that the CCV dynamically interacts with host recycling endosomes for vacuolar expansion and potentially uncovers novel host cell factors essential for *Coxiella* pathogenesis.

## 1 Introduction

*Coxiella burnetii* is a highly infectious, zoonotic, bacterial pathogen and the causative agent of human Q (query) fever. In acute cases, Q fever manifests as a flu-like illness, whereas in about 5% of patients, the disease progresses to a chronic stage that often leads to life-threatening endocarditis [1]. The treatment plan for chronic Q fever involves an 18-month-long combined antibiotic therapy and currently, there is no Q fever vaccine licensed in the U.S., presenting challenges in both treating and preventing the disease. Q fever has emerged as a significant public health concern worldwide [2-10].

In the course of a natural infection, *Coxiella* is transmitted through the respiratory route, where alveolar macrophages within the lung parenchyma endocytose the bacteria [11, 12]. Following entry into host cells, *Coxiella* resides in a tight-fitting phagosome that expands and matures over the next 48-72 hours by interacting with the host endocytic pathway [13]. The phagosome maturation pathway involves sequential fusion with the host early and late endosomes, incorporating their respective membrane markers Rab5 and Rab7 [13, 14] and subsequent fusion with lysosomes, acquiring lysosomal markers such as LAMP1 (lysosome-associated membrane glycoprotein-1) and vacuolar ATPase [15-19]. The phagosome-lysosome fusion is a unique event in the *Coxiella* life cycle that generates an acidic phagolysosome-like vacuole with a pH of ∼5.2 [20, 21], within which the bacteria multiply. The mature phagosome, termed the *Coxiella-*containing vacuole (CCV), is crucial to *Coxiella* metabolism, replication, and pathogenesis [14, 22, 23].

Previously, using a ratiometric fluorescence-based pH measurement assay, we demonstrated that although the CCV is acidic, further escalation in CCV acidity is bactericidal to *Coxiella* [20] and therefore, *Coxiella* actively regulates CCV acidity in a type 4B secretion system (T4BSS)-dependent manner [21]. Specifically, we showed that *Coxiella* T4BSS inhibits the host endosomal maturation pathway, markedly reducing the number of acidic, mature endosomes and lysosomes available for fusion with the CCV [21]. Since the CCV attains its luminal acidity by fusing with mature endosomes and lysosomes, inhibiting endosomal maturation indirectly aids *Coxiella* in regulating the CCV pH [21]. One notable outcome of *Coxiella* inhibition of endosomal maturation is the increase in mean endosomal pH (pH ∼5.2) in the infected cells. Since mature endosomes have a significantly lower pH (∼ 4.7), we define the endosomes with an elevated pH as ‘immature’ endosomes. Because endosomal maturation is crucial to cargo transport and degradation [24], the ‘immature’ endosomes in *Coxiella-*infected cells may be impaired in these functions and therefore detrimental to host cells. However, the fate and the trafficking pattern of the ‘immature’ endosomes in *Coxiella*-infected cells remained unclear. In this study, we aim to characterize the endosomal trafficking in *Coxiella-*infected cells by using HeLa cells expressing endosome-specific fluorescent markers analyzed by quantitative confocal microscopy.

The Ras-associated binding (Rab) proteins are the largest family in the Ras superfamily GTPases and are the master regulators of all vesicular trafficking in mammalian cells [25]. Rab proteins localize on the cytosolic surface of endosomes and perform functions like cargo selection and sorting, vesicle transport along microtubules, and vesicle tethering and fusion with target organelles [25]. The human genome encodes more than 60 Rab proteins that are associated with specific endosome populations to regulate their trafficking, sorting, and recycling events. Rab proteins cycle between an active, GTP-bound, membrane-recruited form and an inactive, GDP-bound, cytosolic form to associate and dissociate with specific endosomes [24, 25]. Therefore, the localization of a specific Rab protein provides key information about the trafficking pattern of a population of endosomes in mammalian cells.

To understand the trafficking of the *Coxiella-*generated ‘immature’ endosomes, we ectopically expressed EGFP-tagged Rab proteins in the infected host cells and analyzed their localization by confocal microscopy. To our surprise, we observed that the CCV, in addition to acquiring Rab5 and Rab7, acquired the recycling endosomal markers Rab11a and Rab35, suggesting that the CCV also interacts with the host recycling endosomes during maturation. Furthermore, we observed an increase in Rab11a and Rab35-positive endosomes in the infected cells, indicating that *Coxiella* enhances the recycling endosome content in the infected cells. Finally, silencing Rab11a and Rab35 using siRNA resulted in significantly smaller CCVs, suggesting that recycling endosomes facilitate CCV expansion. Our study is the first report suggesting a CCV-host recycling endosome interaction indicating the importance of recycling endosomes in the *Coxiella* intracellular life cycle.

## 2 Materials and methods

### 2.1 Bacteria and mammalian cells

*Coxiella burnetii* Nine Mile Phase II (NMII) (Clone 4, RSA 439) and mCherry-expressing *C. burnetii* NMII [26] were grown and stored as previously described [21]. Human cervical epithelial cells (HeLa, ATCC CCL-2) were maintained in Roswell Park Memorial Institute (RPMI) 1640 medium with glutamine (Cat. 10-040-CV, Corning, New York, NY) and 10% fetal bovine serum (FBS; Cat. S11150, R&D Systems, Inc., Minneapolis, MN). The genome equivalents (GE) for the *Coxiella* stocks were determined by qPCR using primers for the *Coxiella burnetii dotA* gene [27]. The multiplicities of infection (MOI) of the bacteria stocks were optimized for HeLa cells and each type of culture vessels to ∼1 internalized bacterium per cell at 37°C and 5% CO_2_.

### 2.2 Preparation of EGFP-tagged Rab plasmids

The plasmids encoding EGFP-tagged Rab proteins were generous gifts from Dr. Mary Weber, University of Iowa [28]. The plasmids were first transformed into competent *E. coli* DH5α (Cat. C2988J, New England Biolabs, Ipswich, MA) and stored at -80°C. A single colony of the transformed *E. coli* was grown in 5 mL lysogeny broth (LB) containing 50 μg/mL kanamycin or 100 μg/mL carbenicillin overnight, which was then used to inoculate a 150 mL LB culture and incubated overnight. The GeneJET Endo-free Plasmid Maxiprep Kit (Cat. K0861, Thermo Fisher Scientific Baltics UAB, Lithuania) was used to isolate and purify plasmid DNA from bacterial cultures according to the manufacturer’s protocol. Plasmids were then concentrated using the MilliporeSigma Amicon Ultra-0.5 Centrifugal Filter Units (Cat. UFC503096, Fisher Scientific, Waltham, MA) and stored at -20°C until used for cell transfection.

### 2.3 Transfection and immunofluorescent assay

HeLa cells, grown in RPMI with 10% FBS (hereafter referred to as 10% RPMI), were plated in a 24-well plate (2×10^4^ cells per well) and simultaneously transfected with 0.4 μg of respective EGFP-Rab plasmids using FuGENE 6 transfection reagent (Cat. E269A, Promega, Madison, WI) per the manufacturer’s protocol. At 24 hours post-transfection, the cells were infected with mCherry-*Coxiella* in 250 uL 10% RPMI for 1 h, washed extensively with sterile phosphate-buffered saline (PBS; HyClone, Cat. SH30256.01, Cytiva, Marlborough, MA), and incubated in 10% RPMI at 37°C. On the day before the indicated time points, cells were trypsinized, counted, and diluted to 1×10^5^ cells/mL. Diluted cells were then plated onto coverslips placed in a separate 24-well plate (5×10^4^ cells per coverslip) and allowed to adhere overnight. The next day, cells were fixed in 2.5% paraformaldehyde (PFA; Cat. 15710, Electron Microscopy Sciences, Hatfield, PA) for 15 min, washed in PBS, and blocked/permeabilized for 20 min in 1% bovine serum albumin (BSA; Cat. BP9700-100, Fisher BioReagents, Pittsburgh, PA) and 0.1% saponin (Cat. S0019, TCI America, Portland, OR) in PBS. Cells were then incubated in rabbit anti-LAMP1 (1:1000; Cat. PA1-654A, Thermo Fisher Scientific, Waltham, MA) primary antibody for 1h followed by Alexa Fluor secondary antibody (1:1000; Life Technologies) for 1 h to label the CCV. Following washing with PBS, coverslips were mounted using ProLong Gold Antifade Mountant with 4’, 6’-diamidino-2 phenylindole (DAPI; Cat. P36941, Invitrogen, Carlsbad, CA) and visualized on a Leica Stellaris confocal microscope using a 63x oil immersion objective. Images were processed and analyzed in ImageJ FIJI [29].

### 2.4 Enumeration of Rab11a/Rab35-positive CCVs

HeLa cells were plated in a 24-well plate and transfected with plasmids encoding EGFP-tagged Rab11a or Rab35 using FuGENE 6 transfection reagent according to the manufacturer’s protocol. At 24 hours post-transfection, the cells were infected with mCherry-*Coxiella* in 250 μL 10% RPMI for 1 h, washed in PBS, and incubated in 10% RPMI. On the day before the indicated time points, cells were trypsinized, diluted to 1×10^5^ cells/mL, and plated onto coverslips in a 24-well plate (5×10^4^ cells per coverslip). The next day, the cells were subjected to immunofluorescent assay (IFA) where they were fixed with 2.5% PFA for 15 min and blocked/permeabilized with 1% BSA in PBS with 0.1% saponin for 20 min. Cells were then incubated in rabbit anti-LAMP1 (1:1000; Cat. PA1-654A, Thermo Fisher) primary antibody for 1 h followed by Alexa Fluor 647 goat anti-rabbit secondary antibody (1:1000; Cat. A21245, Invitrogen) for 1 h to label the CCVs. Following washing with PBS, the coverslips were mounted with ProLong Gold Antifade Mountant with DAPI. The cells were visualized with a Leica Stellaris confocal microscope using a 63x oil immersion objective and images were imported and analyzed in ImageJ FIJI. A Rab-LAMP1 colocalization at the CCV membrane was considered positive for Rab trafficking to the CCV. Approximately 100 CCVs were scored for Rab11a and Rab35 localization and the percentage of Rab11a- and Rab35-positive CCVs were determined. The statistical difference in the percentage of Rab11a and Rab35-positive CCVs at each time-point was determined by Welch’s t-test from three independent experiments (N=3).

### 2.5 Quantification of Rab11a and Rab35 content in *Coxiella-*infected cells

HeLa cells were either mock-infected or infected with mCherry-*Coxiella* in 24-well plates (2.5×10^4^ cells per well; two wells per condition) for 1 h, washed extensively with PBS, and incubated in 10% RPMI. For 3-day experiments, the cells were trypsinized at 2 dpi, diluted to 1×10^5^ cells/mL, and replated onto coverslips in a new 24-well plate (5×10^4^ cells per coverslip). At 3 dpi, cells were fixed with 2.5% PFA for 15 min and washed thoroughly with PBS. For 6-day experiments, cells were trypsinized, diluted, and replated first at 3 dpi and again at 5 dpi (5×10^4^ cells per coverslip). Next day, cells were fixed with 2.5% PFA and washed thoroughly with PBS. For both 3-day and 6-day experiments, following fixation, the cells were blocked/permeabilized with 1% BSA in PBS with 0.1% saponin for 20 min. Cells were then incubated overnight at 4°C in rabbit anti-Rab35 (1:100; Cat. 11329-2-AP, Proteintech, Rosemont, IL) or for 1 h at room temperature (RT) in rabbit anti-Rab11a (1:500; Cat. 20229-1-AP, Proteintech) primary antibody. The coverslips were washed with PBS and then incubated in Alexa Fluor 488 goat anti-rabbit (1:1000; Cat. A11034, Invitrogen) secondary antibody for 1 h at RT. Upon washing with PBS, the coverslips were mounted using ProLong Gold with DAPI. Cells were visualized with a Leica Stellaris confocal microscope using a 63x oil immersion objective. To quantify Rab content from mock and WT-infected cells, identically captured and processed images were imported into ImageJ FIJI and the total fluorescent intensity of Rab11a or Rab35 was measured and normalized to cell area. At least 25 cells were analyzed per condition in each of three independent experiments (N=3). The statistical difference in Rab11a and Rab35 intensities between mock and *Coxiella*-infected cells was measured by Welch’s t-test.

### 2.6 RNA interference and immunoblotting

HeLa cells (10^5^ cells per well, 6-well plate) were reverse transfected with 50 nM small-interfering RNA (siRNA) SMARTpools specific for human Rab11a (ON-TARGETplus, Cat. L-004726-00-0005, Dharmacon Inc., Horizon Discovery Ltd., Lafayette, CO), Rab35 (ON-TARGETplus, Cat. L-009781-00-0005, Dharmacon Inc.), or a non-targeting (NT) control (Cat. D-001810-10-05, Dharmacon Inc.) using DharmaFECT-1 transfection reagent (Cat. T-2001-01, Dharmacon Inc.) in RPMI with 5% FBS according to the manufacturer’s protocol. At 48 h post-transfection, cells were infected with mCherry-*Coxiella* for 1 h at 37°C. Following washing with PBS, cells were harvested by trypsinization and subjected to a second round of siRNA transfection in a 24-well plate (2.5×10^4^ cells per well). On the day before the indicated time points, cells were trypsinized, resuspended to 1×10^5^ cells/mL, and either replated onto coverslips (2.5×10^4^ cells per coverslip) or into another 24-well plate without coverslips. The next day, the cells on coverslips were fixed with 2.5% PFA and subjected to immunofluorescent assay with rabbit anti-LAMP1 primary and Alexa Fluor 488 secondary antibodies to label the CCVs. Concurrently, mock-infected cells from the second 24-well plate were lysed with RIPA lysis buffer (Cat. 89900, Thermo Fisher) supplemented with HALT protease inhibitor (Cat. 87786, Thermo Fisher) on ice. The soluble fraction from the cell lysates were collected and protein concentrations were determined using the Pierce Rapid Gold BCA kit (Cat. A53226, Thermo Fisher). To confirm Rab11a and Rab35 silencing, 10 μg of total protein for both NT and Rab siRNA-transfected samples were resolved in 4-20% gradient SDS-PAGE (Mini-PROTEAN Gel, Cat. 4561094, Bio-Rad Laboratories, Hercules, CA) and subsequently transferred to a PVDF membrane for immunoblotting (Immun-Blot PVDF Membrane, Cat. 1620174, Bio-Rad). After blocking in Bio-Rad EveryBlot buffer (Cat. 12010020, Bio-Rad) for 5 min, the membranes were then probed separately using rabbit anti-Rab11a (Cat. 20229-1-AP, Proteintech), rabbit anti-Rab35 (Cat. 11329-2-AP, Proteintech), and anti-β-tubulin (BT7R) (HRP-conjugated loading control; Cat. MA5-16308-HRP, Invitrogen) antibodies in 1% BSA in PBS. After washing, the membranes were incubated with horseradish peroxidase-conjugated goat anti-rabbit (1:1000; Cat. 31460, Thermo Fisher) secondary antibody in EveryBlot blocking buffer, and developed using enhanced chemiluminescence (ECL) reagent (Cat. 170-5060, Bio-Rad).

### 2.7 Measurement of CCV area

HeLa cells were reverse-transfected with NT control, Rab11a, or Rab35-specific siRNAs as described above, and at 48 h post-transfection, were infected with mCherry-*Coxiella* for 1 h. Upon washing with PBS, cells were subjected to a second round of siRNA transfection as described above. On the day before the indicated time points, cells were trypsinized, resuspended to 1×10^5^ cells/mL, and replated onto coverslips (2.5×10^4^ cells/coverslip). The next day, the coverslips were fixed with 2.5% PFA and cells were immunostained with rabbit anti-LAMP1 primary antibody and Alexa Fluor secondary antibody to label the CCVs. Mounted coverslips were visualized using a Keyence BZ-X710 fluorescent microscope. Images of individual CCVs were captured and processed identically between conditions, and the CCV areas from exported images were measured in ImageJ FIJI. A 20 μm scale bar placed on a representative image for each sample was used for setting the scale in ImageJ. CCV areas were expressed as pixels and plotted using GraphPad Prism. At least 25 CCVs were measured per condition for each of the three independent experiments (N=3). The statistical differences in CCV areas between non-targeting and Rab11a/Rab35-specific siRNA-transfected cells were measured by Welch’s t-test.

### 2.8 Data analyses

Fluorescent and confocal images were processed and analyzed in ImageJ FIJI software [29]. All statistical analyses were performed using unpaired t-test with Welch’s correction (Welch’s t-test) using GraphPad Prism (GraphPad, La Jolla, CA). A p-value < 0.05 was considered significant. Data are plotted as mean ± standard error of mean (SEM).

## 3 Results

### 3.1 Recycling endosomal markers Rab11a and Rab35 localize to the CCV during infection

We previously observed that *Coxiella* actively regulates the CCV acidity by inhibiting host endosomal maturation in a T4BSS-dependent manner [21]. However, by impeding endosomal maturation, *Coxiella* gives rise to population of ‘immature’ endosomes with a higher mean pH compared to mature endosomes. As the immature endosomes may be impaired in cargo trafficking and degradation, they may be detrimental to the host cells. However, whether *Coxiella* manipulates the trafficking of the ‘immature’ endosomes in the infected cells has not been clear. Since Rab proteins are the master regulators of endocytic trafficking, we hypothesized that *Coxiella* manipulates one or more Rab proteins to modulate endosomal trafficking in the infected cells. To test this, we screened 13 major Rab proteins in *Coxiella-*infected cells for their interaction with the CCV. Plasmids encoding EGFP-tagged Rab proteins (Table 1; a gift from Dr. Mary Weber, University of Iowa) were maxiprepped from *E. coli* culture and stored in endotoxin free elution buffer. HeLa cells were transfected with the EGFP-Rab plasmids and infected with mCherry-*Coxiella*. At 3 dpi, the cells were immunostained with LAMP1 primary antibody to label the CCVs. The cells were then visualized with a Leica Stellaris confocal microscope where 488 and 647 nm lasers were used for imaging the Rab proteins and LAMP1, respectively. Images were processed and analyzed in ImageJ FIJI and a Rab-LAMP1 colocalization at the CCV was considered positive for Rab trafficking to the CCV. EGFP tagged-Rab5 and Rab7 were included as positive controls in the screen as Rab5 and Rab7 are known to localize to the CCV during infection [30, 31]. As expected, all CCVs were positive for Rab5, Rab7, and LAMP1. In addition to that, we interestingly observed that some CCVs were also positive for Rab11a and Rab35 as evidenced by their colocalization with LAMP1 at the CCV membrane (Figure 1). Both Rab11a and Rab35 are associated with host recycling endosomes, where they regulate the ‘slow’ and ‘fast’ endosomal recycling pathways, respectively [32-34]. Therefore, these data suggest that, in addition to interacting with early endosomes, late endosomes, and lysosomes, at least a subset of CCVs also interacts with the host recycling endosomes in both the ‘slow’ and the ‘fast’ recycling pathways. A CCV area comparison between the Rab11a/Rab35-positive and the negative CCVs showed no significant difference in areas between the positive and the negative CCVs (data not shown). Moreover, *Coxiella* colonies appeared unaffected by the presence of Rab11a and Rab35 at the CCV. This study is the first report suggesting that the CCV interacts with host recycling endosomes and acquires recycling endosomal markers during maturation.

**Table 1.**
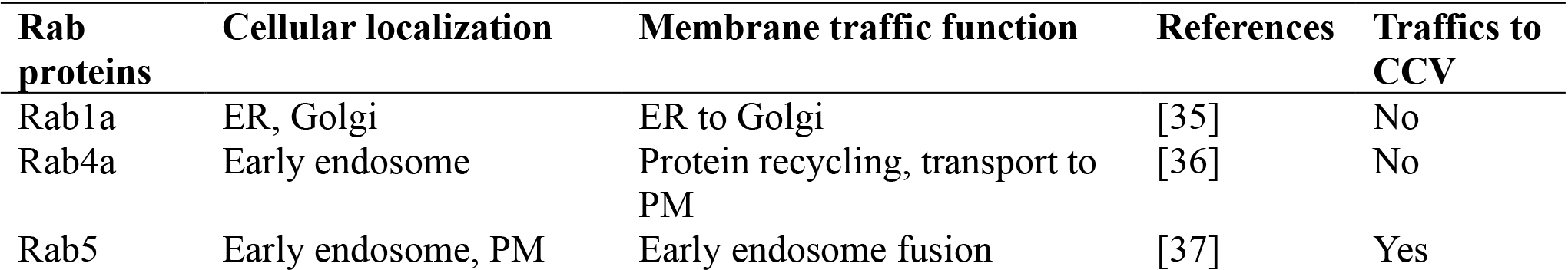
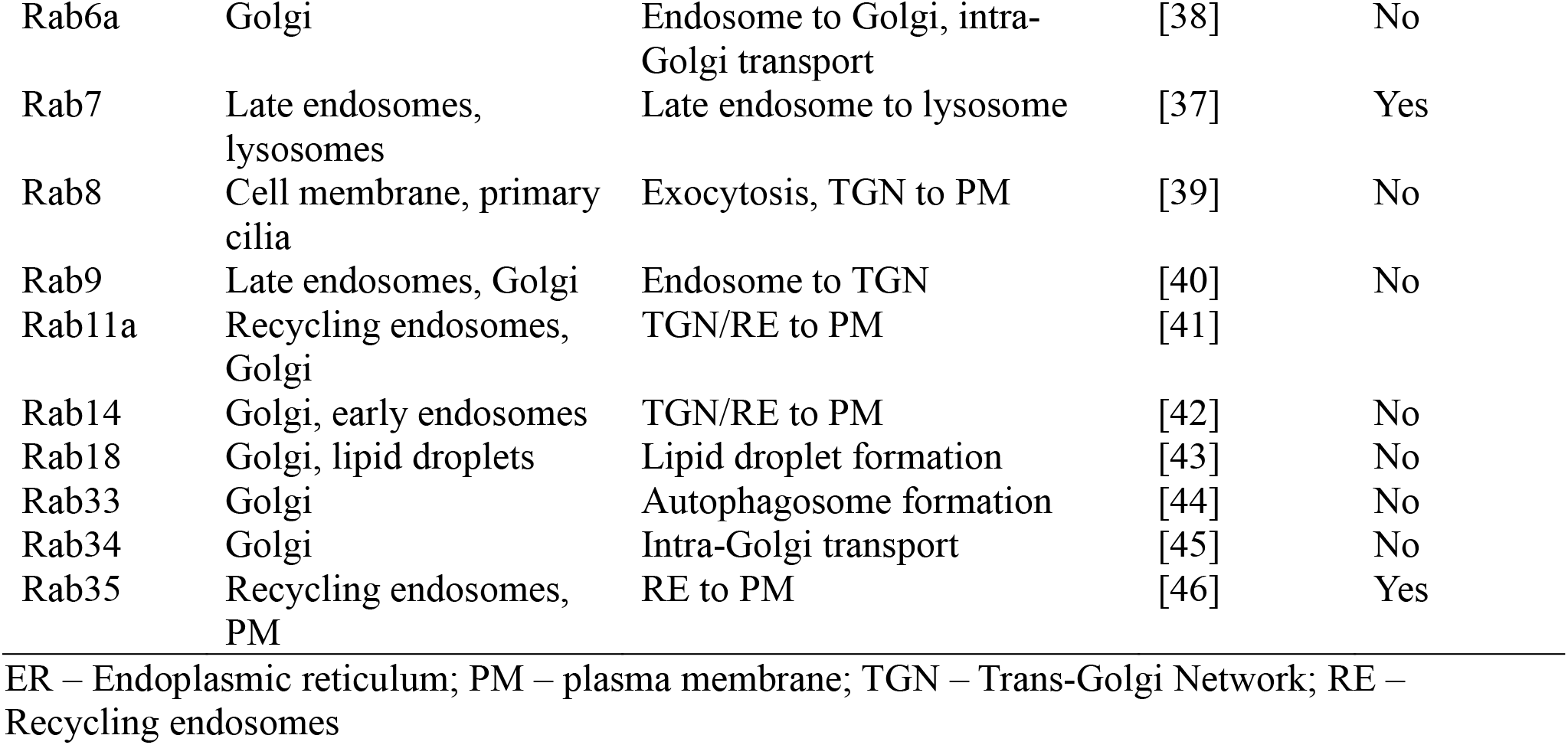
EGFP-tagged Rab proteins used in this study.

| Rab proteins | Cellular localization | Membrane traffic function | References | Traffics to CCV |
| --- | --- | --- | --- | --- |
| Rab1a | ER, Golgi | ER to Golgi | [35] | No |
| Rab4a | Early endosome | Protein recycling, transport to PM | [36] | No |
| Rab5 | Early endosome, PM | Early endosome fusion | [37] | Yes |
| Rab6a | Golgi | Endosome to Golgi, intra-Golgi transport | [38] | No |
| Rab7 | Late endosomes, lysosomes | Late endosome to lysosome | [37] | Yes |
| Rab8 | Cell membrane, primary cilia | Exocytosis, TGN to PM | [39] | No |
| Rab9 | Late endosomes, Golgi | Endosome to TGN | [40] | No |
| Rab11a | Recycling endosomes, Golgi | TGN/RE to PM | [41] |  |
| Rab14 | Golgi, early endosomes | TGN/RE to PM | [42] | No |
| Rab18 | Golgi, lipid droplets | Lipid droplet formation | [43] | No |
| Rab33 | Golgi | Autophagosome formation | [44] | No |
| Rab34 | Golgi | Intra-Golgi transport | [45] | No |
| Rab35 | Recycling endosomes, PM | RE to PM | [46] | Yes |

ER – Endoplasmic reticulum; PM – plasma membrane; TGN – Trans-Golgi Network; RE – Recycling endosomes

**Figure 1.**
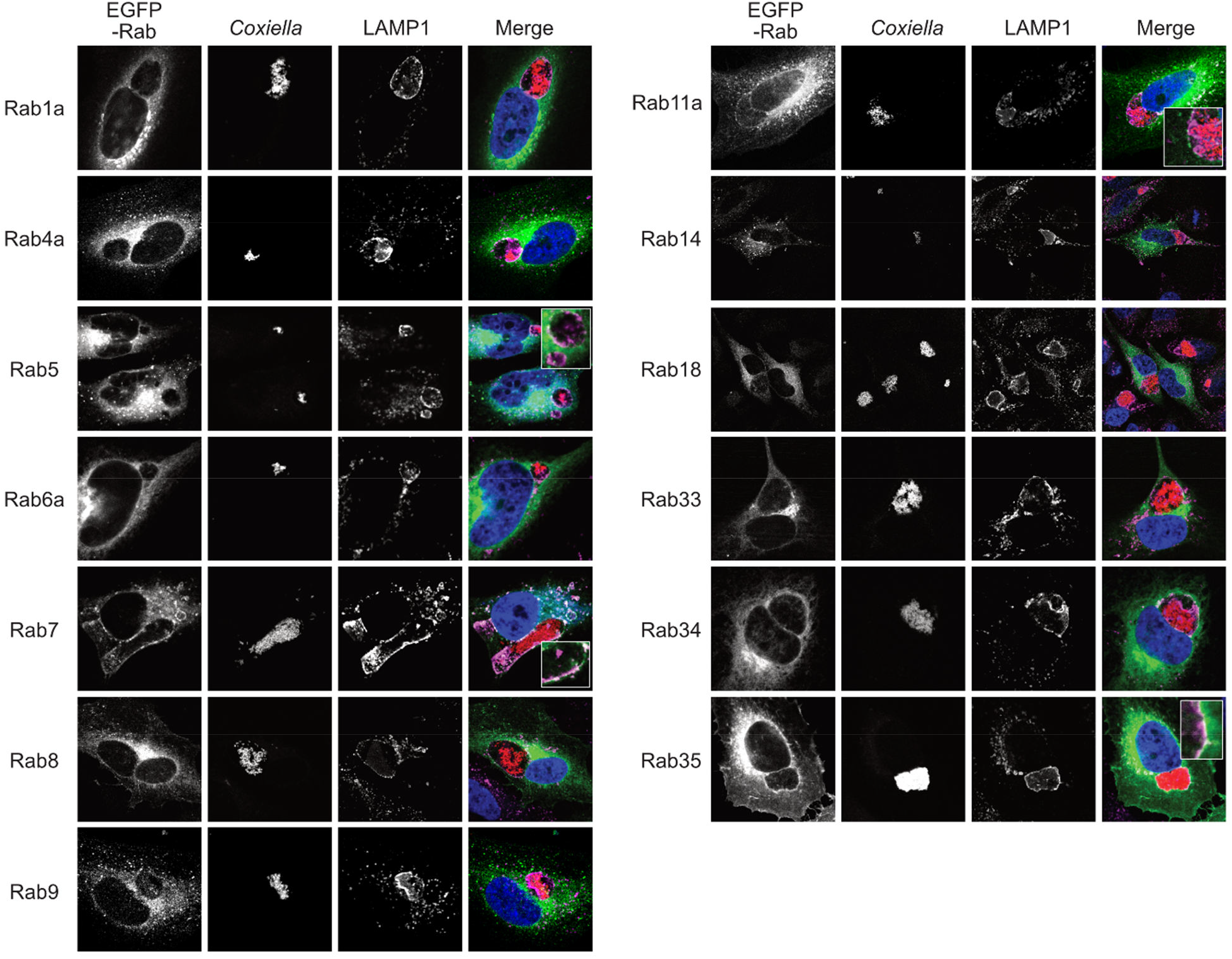
Host recycling endosome-associated proteins Rab11a and Rab35 localize to the CCV. Representative images of *Coxiella-*infected HeLa cells, expressing EGFP-tagged Rab proteins. HeLa cells were transfected with plasmids encoding EGFP-Rab proteins and at 24 h post-transfection, were infected with mCherry-*Coxiella* for 1 h. At 3 dpi, cells plated on coverslips were fixed and subjected to immunofluorescent assay with anti-LAMP1 antibody to label the CCVs. Cells were visualized with a Leica Stellaris confocal microscope and images were analyzed in ImageJ FIJI. CCVs, where LAMP1 and Rab proteins colocalized at the membrane (shown in insets) were considered positive for a specific Rab protein. As expected, CCVs were positive for both Rab5 and Rab7 at 3 dpi. Additionally, we observed that Rab11a and Rab35 also localized to CCVs. Notably, only a subset of CCVs were positive for Rab11a or Rab35.

### 3.2 CCV-Rab11a and CCV-Rab35 interactions are dynamic and temporally regulated

In our Rab localization screen, we interestingly observed that only a subset of the CCVs were positive for Rab11a or Rab35 at a given time-point. This led us to hypothesize that the interaction between the CCV and Rab11a/Rab35 are dynamic in nature. To test this, we quantified the percentage of Rab11a- and Rab35-positive CCVs in a population of *Coxiella-*infected HeLa cells at 3 and 6 dpi. Rab11a- or Rab35-positive and negative CCVs were identified in ImageJ FIJI by a colocalization of Rab and LAMP1 at the CCV membrane, as described above (Figure 2A). Our quantitation data revealed that at 3 dpi, ∼52% of CCVs were positive for Rab11a, whereas only ∼39% of CCVs were positive for Rab35 (Fig. 2B-C). Interestingly this pattern reversed at 6 dpi, when only ∼22% of CCVs were positive for Rab11 and ∼64% of CCVs were positive for Rab35 (Fig. 2B-C). Together, these data suggest that, while CCVs preferentially interact with the Rab11a-positive endosomes at 3pi, at 6 dpi, the preferential interaction shifts towards the Rab35-positive endosomes. Therefore, our data indicate that Rab11a and Rab35 interaction with the CCV are dynamic and varies depending on the stage of infection.

**Figure 2.**
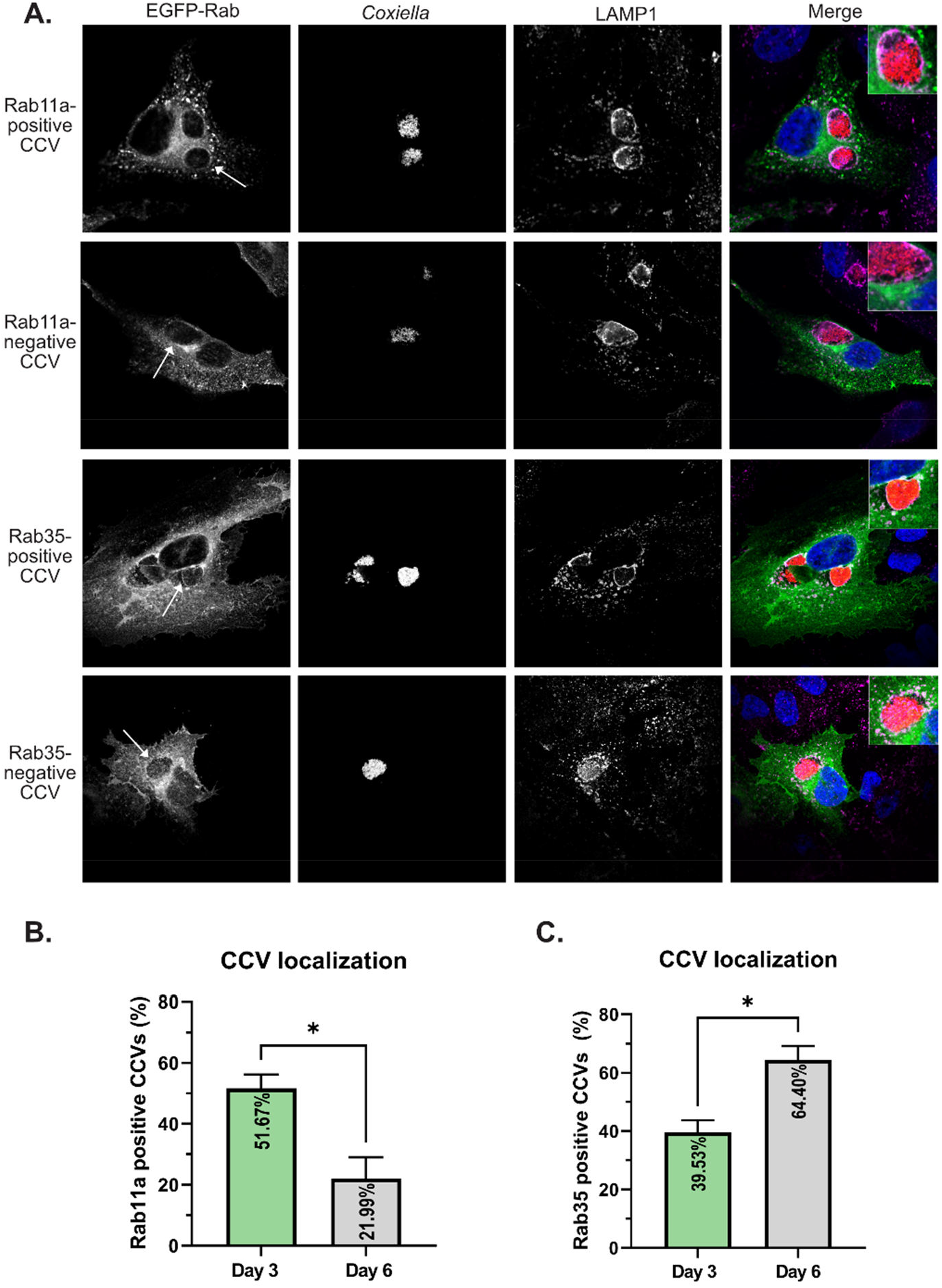
CCV-Rab11a and CCV-Rab35 interactions are dynamic and temporally regulated. **(A)** Representative images showing Rab11a and Rab35-positive and negative CCVs. HeLa cells were transfected with plasmids encoding EGFP-Rab11a or EGFP-Rab35 and at 24 h post-transfection, were infected with mCherry-*Coxiella* for 1 h. At 3 dpi, cells plated on coverslips were fixed and subjected to immunofluorescent assay with anti-LAMP1 antibody to label the CCVs. Cells were visualized with a Leica Stellaris confocal microscope and images were analyzed in ImageJ FIJI. CCVs, where LAMP1 and Rab proteins colocalized at the membrane (shown in insets) were considered positive for a specific Rab protein. CCVs were scored for Rab11a or Rab35 localization and percentage of positive CCVs were calculated. **(B-C)** Quantification of Rab11a- and Rab35-positive and negative CCVs at 3 and 6 dpi revealed that Rab11a localized to significantly more CCVs at 3 dpi compared to 6 dpi, whereas the pattern reversed for Rab35, which localized to more CCVs at 6 dpi. Data shown as mean±SEM of at least 100 infected cells per condition in each of three independent experiments (N=3) as analyzed by Welch’s t-test; *, P<0.05.

### 3.3 *Coxiella* infection leads to increased recycling endosome content in infected cells

Our observation that *Coxiella* interacts with two different populations of recycling endosomes in the host cells led us to hypothesize that *Coxiella* manipulates host recycling endosomes during infection. To test this hypothesis, we quantitated the total cellular Rab11a and Rab35 content in mock and *Coxiella-*infected HeLa cells by immunofluorescent assay with anti-Rab11a or anti-Rab35 antibody. EGFP-Rab11 or EGFP-Rab35 transfected HeLa cells were not used in this experiment because of potential differences in ectopic expression level of Rab proteins between individual cells. Mock and *Coxiella-*infected HeLa cells were immunostained using anti-Rab11a or anti-Rab35 antibodies at 3 and 6 dpi and identically captured images were analyzed in ImageJ FIJI. The total fluorescent intensity of Rab11 or Rab35 was measured from mock and *Coxiella-*infected cells normalized to cell area and were considered indicative of total recycling endosome content in each cell. Our data revealed a ∼19% increase in Rab11a fluorescent intensity (p < 0.0001; Figures 3A, B) and a ∼27% increase in Rab35 fluorescent intensity in infected cells compared to mock at 3 dpi (p < 0.0001; Figures 4A, B). However, to our surprise, we did not observe any difference in Rab11a or Rab35 fluorescent intensities between mock and *Coxiella*-infected cells at 6 dpi (Figures 3C, 4C). These data suggest that *Coxiella* increases the recycling endosome content in infected host cells at 3 dpi, but not at 6 dpi.

**Figure 3.**
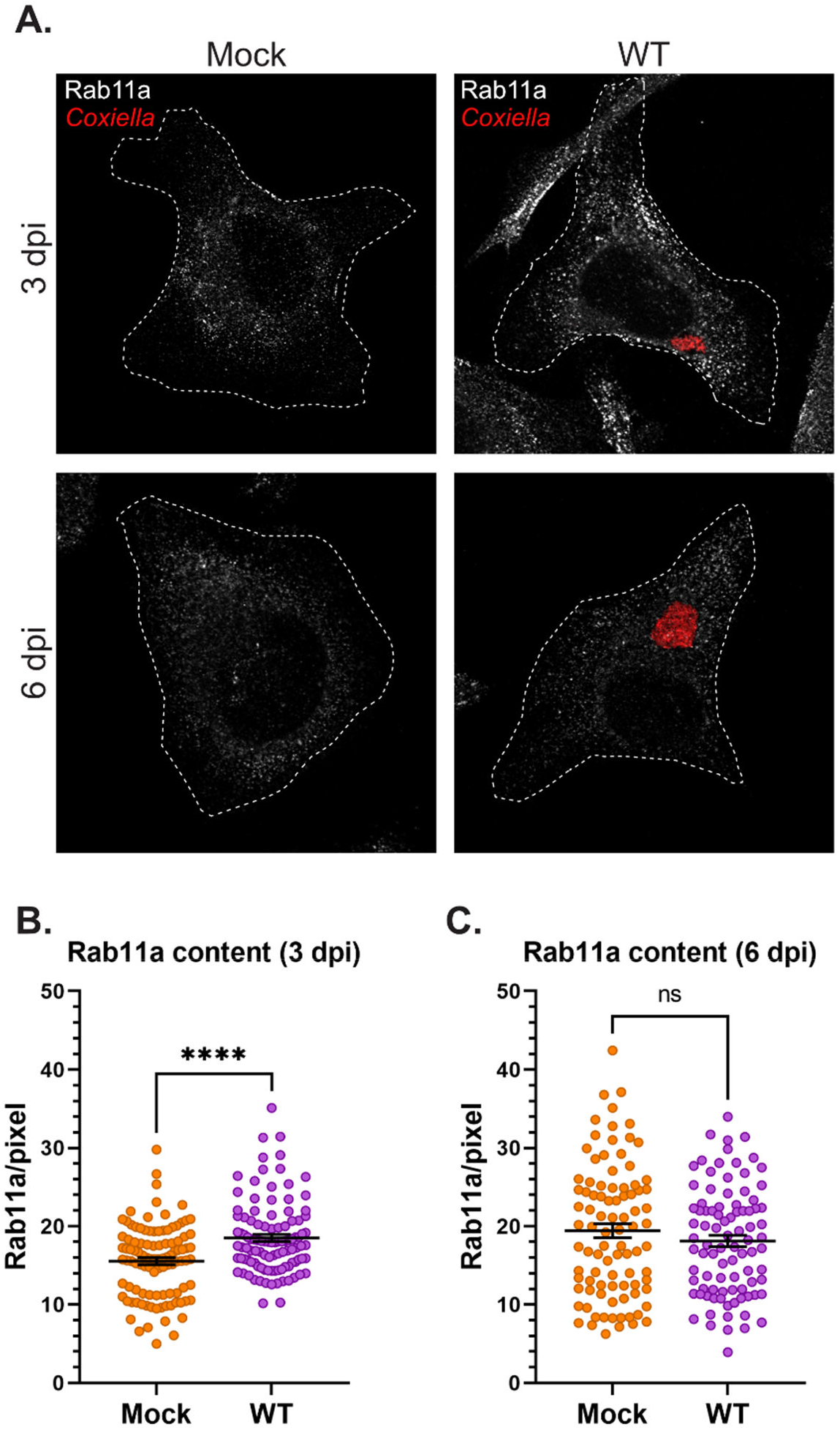
*Coxiella* increases Rab11a content in infected host cells. **(A)** Representative images showing Rab11a-positive recycling endosome content of mock and *Coxiella-*infected HeLa cells at 3 and 6 dpi. HeLa cells were mock infected or infected with mCherry-*Coxiella* for 1 h. On the days before the indicated time points, the cells were plated on coverslips. The next day, cells were fixed and subjected to immunofluorescent assay with anti-Rab11a antibody to uniformly stain the Rab11a-positive recycling endosomes. Cells were visualized with a Leica Stellaris confocal microscope and the images were analyzed in ImageJ FIJI. Identically captured images were imported into ImageJ FIJI and the total fluorescent intensity of Rab11a was measured and normalized to cell area. **(B-C)** Quantification of Rab11a fluorescent intensity revealed an increase in Rab11a content in *Coxiella-* infected cells at 3 dpi but not at 6 dpi. Data shown as mean±SEM of at least 25 cells per condition in each of three independent experiments (N=3) as analyzed by Welch’s t-test; ****, P<0.0001; ns, non-significant.

**Figure 4.**
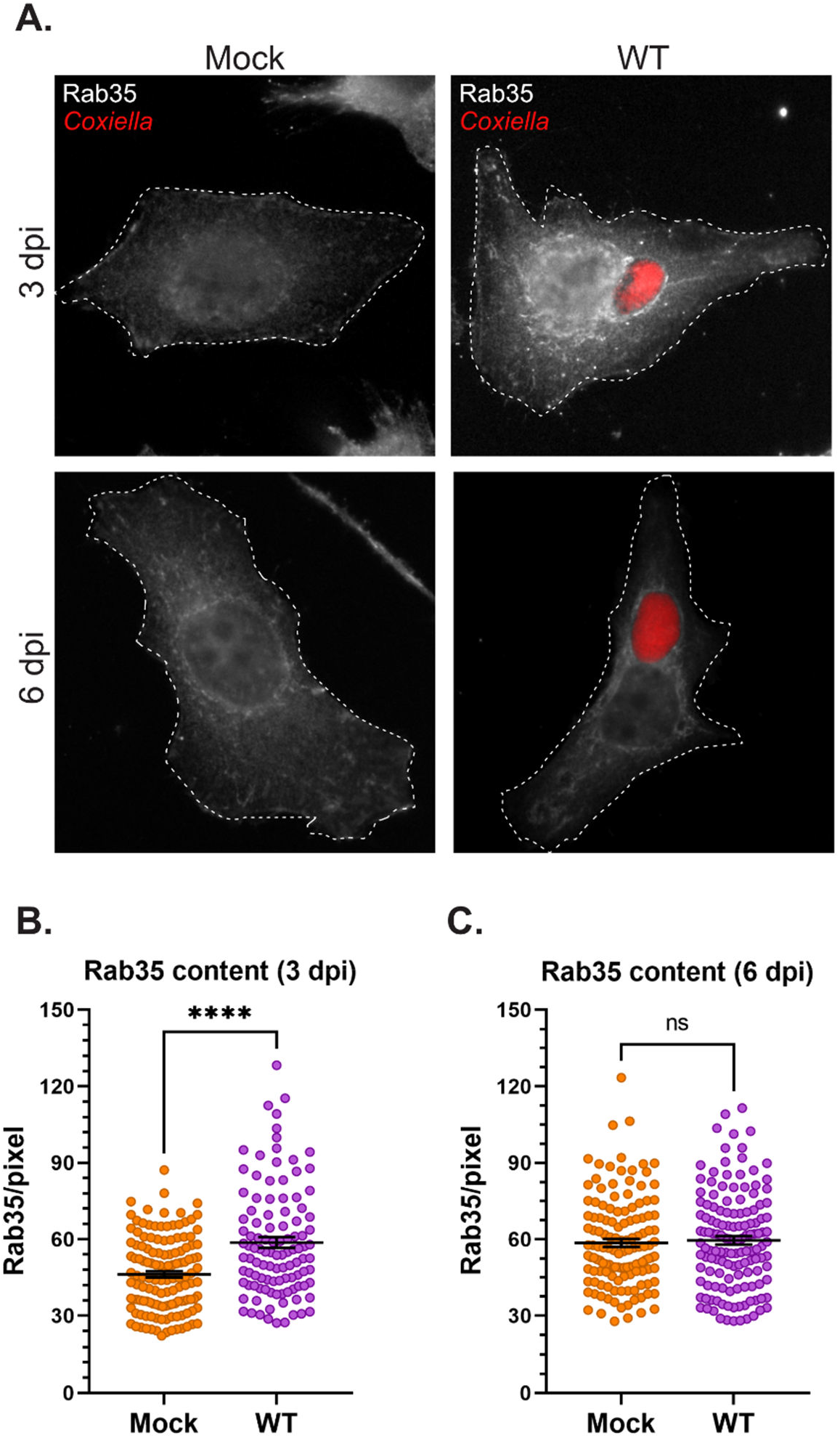
*Coxiella* increases Rab35 content in infected host cells. **(A)** Representative images showing Rab35-positive recycling endosome content of mock and *Coxiella-*infected HeLa cells at 3 and 6 dpi. HeLa cells were mock infected or infected with mCherry-*Coxiella* for 1 h. On the days before the indicated time points, the cells were plated on coverslips. The next day, cells were fixed and subjected to an immunofluorescent assay with anti-Rab35 antibody to uniformly stain the Rab35-positive recycling endosomes. Cells were visualized with a Leica Stellaris confocal microscope and the images were analyzed in ImageJ FIJI. Identically captured images were imported into ImageJ FIJI and the total fluorescent intensity of Rab35 was measured and normalized to cell area. **(B-C)** Quantification of Rab35 fluorescent intensity revealed an increase in Rab35 content in *Coxiella-* infected cells at 3 dpi but not at 6 dpi. Data shown as mean±SEM of at least 25 cells per condition in each of three independent experiments (N=3) as analyzed by Welch’s t-test; ****, P<0.0001; ns, non-significant.

### 3.4 Rab11a and Rab35 silencing inhibits CCV expansion

Based on our observation that the CCV dynamically interacts with recycling endosome-associated Rab11a and Rab35, and that *Coxiella* increases the recycling endosome content of infected cells, we hypothesized that Rab11a and Rab35 positively affects CCV maturation. To test this, we examined CCV expansion in Rab11a or Rab35 knockdown HeLa cells compared to a non-targeting (NT) control using our dual-hit siRNA knockdown protocol described previously (Figure 5A) [21]. Since CCV interaction with Rab11a is higher at 3 dpi, and that with Rab35 is higher at 6 dpi, we measured the CCV areas in Rab11a and Rab35 knockdown cells at 3 dpi and 6 dpi, respectively, to detect maximum effect of the respective knockdowns. Immunoblotting with cell lysates from NT, siRab11a, and siRab35-transfected cells revealed significant depletion of both Rab11a (Figure 5B) and Rab35 (Figure 5C) at 3 and 6 dpi, respectively. The CCV area measurement data revealed a ∼45% reduction in CCV area in Rab11a knockdown cells compared to NT (p<0.0001) at 3 dpi (Figure 5D), and a ∼28% reduction in CCV area in Rab35 knockdown cells (p<0.0001) at 6 dpi (Figure 5E). These data suggest that both Rab11a and Rab35 positively affect CCV expansion at 3 and 6 dpi, respectively, indicating that host recycling endosomes are essential for CCV maturation.

**Figure 5.**
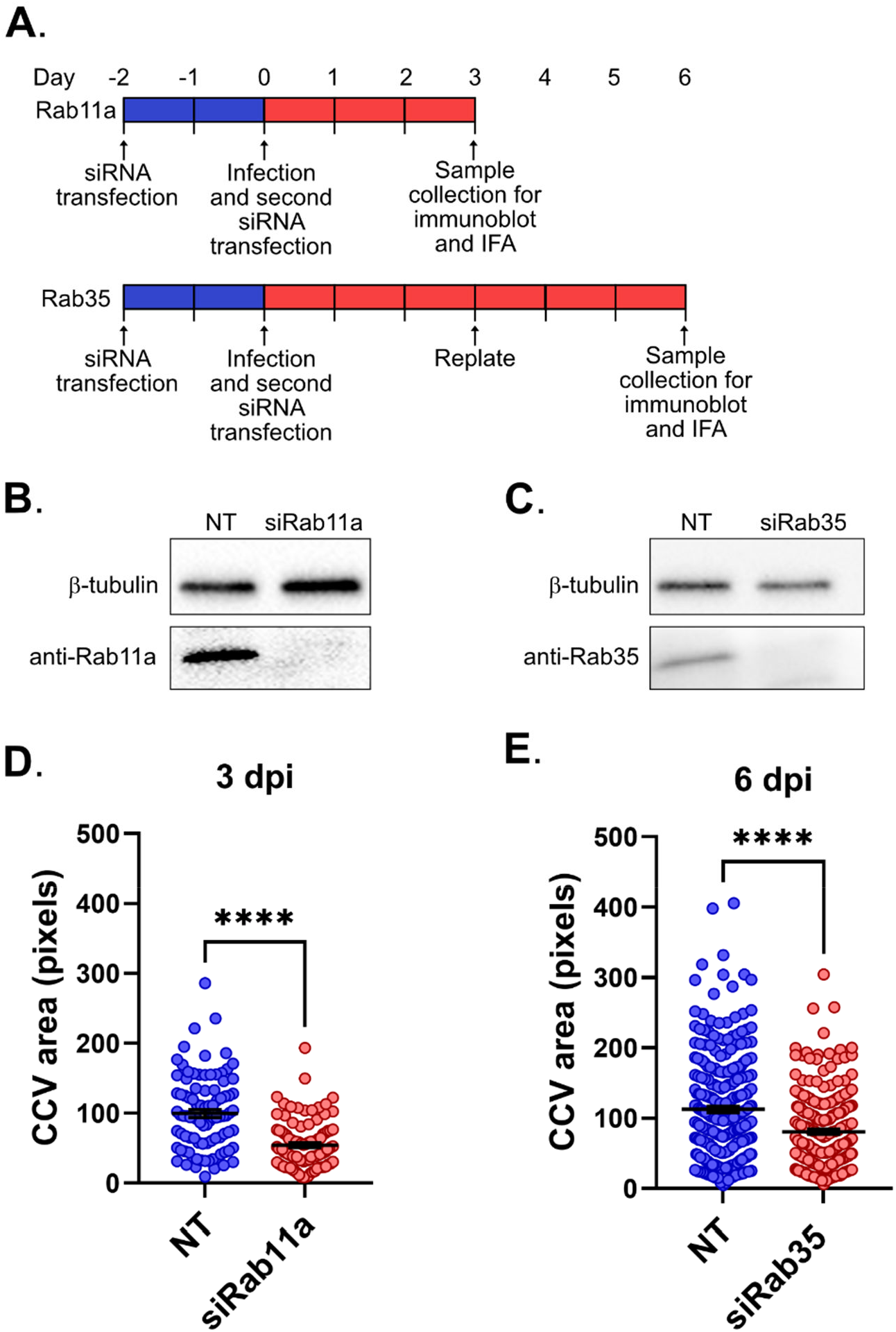
Rab11a and Rab35 are essential for CCV expansion. **(A)** Schematic diagram of dual siRNA transfection protocol in HeLa cells to silence Rab11a and Rab35. Cells were transfected with Rab11a or Rab35-specific siRNA (siRab11a or siRab35, respectively) or a control non-targeting siRNA (NT) and incubated. At 48 h post-transfection cells were infected with mCherry-*Coxiella* and subjected to a second round of siRNA transfection. At the indicated time points, cells were fixed and subjected to immunofluorescent assay with anti-LAMP1 antibody to label the CCVs. **(B-C)** Immunoblots revealed depletion of both Rab11a and Rab35 in siRNA-transfected cells at 3 and 6 dpi, respectively. **(D-E)** measurement of the CCV area revealed significantly smaller CCVs in siRab11a-transfected cells at 3 dpi and siRab35-transfected cells at 6 dpi. Data shown as mean±SEM of at least 25 cells per condition in each of three independent experiments (N=3) as analyzed by Welch’s t-test; ****, P<0.0001; ns, non-significant.

## 4 Discussion

We previously established that *Coxiella* T4BSS inhibits host endosomal maturation for bacterial survival and pathogenesis. However, in the process, *Coxiella* generates a population of ‘immature’ endosomes with an elevated mean pH relative to mature endosomes. To generate a comprehensive understanding of endosomal trafficking in the *Coxiella-*infected cells, we screened 13 well-studied Rab proteins to assess their localization in *Coxiella-*infected cells. We discovered that Rab11a and Rab35, the molecular markers of recycling endosomes, localize to the CCV, although only a subset of CCVs were found positive for Rab11a or Rab35 at a given time point. Our data also revealed that *Coxiella* increases the recycling endosome content of the infected cells at 3 dpi, and Rab11a and Rab35 knockdown resulted in significantly smaller CCVs at 3 and 6 dpi, respectively, suggesting that host recycling endosomes are essential for CCV expansion.

Mammalian endosomal trafficking can be broadly classified into degradative and recycling pathways [47]. While the degradative pathway traffics the cargo through early and late endosomes and finally delivers it to lysosomes for degradation, the recycling pathway sorts and re-exports essential membrane components including receptors, carrier molecules, and cell adhesion molecules that have been internalized [33]. For the first time, we present findings suggesting an interaction between the CCV and the host recycling endosomes. This intriguing observation implies that the CCV not only engages with the host endocytic degradative endosomes but also with recycling endosomes, potentially incorporating their contents and membranes during maturation. Mammalian cells exhibit two simultaneous pathways for cargo recycling namely the ‘fast’ and the ‘slow’ recycling pathways [48]. In the ‘fast’ pathway, the internalized cargo is directly transported back to the plasma membrane from early endosomes. However, the ‘slow’ pathway first traffics the cargo to a perinuclear vesicle known as the endocytic recycling compartment (ERC), within which, the cargo is sorted and then redirected to the plasma membrane via recycling endosomes [33, 49]. Rab35 predominantly regulates the ‘fast’ recycling pathway [32, 34], while Rab11a serves as a major regulator of the ‘slow’ pathway [33, 34, 50]. Therefore, our data suggest that the CCV interacts with both the ‘fast’ and the ‘slow’ recycling pathways during maturation. Moreover, our observation that Rab11a localizes to the CCVs significantly more at 3 dpi whereas Rab35 is more prevalent at CCVs at 6 dpi suggest a preference for interaction with the ‘slow’ recycling pathway during the early stages of maturation and a shift towards a preferential interaction with the ‘fast’ recycling pathway as the CCV matures further.

While the significance of the interactions between CCV and the host recycling endosomes remains uncertain, multiple studies propose that similar interactions with intracellular bacterial vacuoles play a crucial role in providing nutrients to the vacuole or in ensuring vacuolar stability. For example, *Chlamydophila pneumoniae*-containing inclusions recruit the mammalian Rab11/Rab14 adapter protein Fip2, facilitating the early-stage recruitment of Rab11/14 to the inclusion membrane during infection [51, 52]. Rab11/14 positive recycling endosomes trafficked nutrients, such as transferrin to the proximity of *C. pneumoniae* inclusion, although whether transferrin is delivered into the inclusion lumen remained unclear [53-55]. Indeed, a recent study with fluorescent transferrin (Tf488) revealed that in *Coxiella-*infected HeLa cells, Tf488-containing vesicles also trafficked to the proximity of CCVs [56]. Moreover, Rab11-positive recycling endosomes trafficked transferrin in K562 cells [50]. Therefore, it is possible that recycling endosomes traffic transferrin to the CCV aiding in its maturation. Further experimentation is needed to examine this intriguing hypothesis. Another possible benefit of intracellular pathogens subverting recycling endosomes is to augment vacuole stability. For example, *Chlamydia trachomatis* inclusion protein CT229 interacts with Rab35 to maintain the inclusion integrity [28], and *Legionella pneumophila-*containing vacuoles (LCV) acquire Rab11 and Rab35 for stability [57]. It is possible that Rab11 and Rab35 plays a critical role in maintaining CCV stability. However, we did not observe a disintegration of CCV in Rab11a and Rab35 knockdown HeLa cells, raising a possibility that *Coxiella* employes redundant mechanisms to maintain CCV stability.

As mentioned before, we observed that only a subset of CCVs are positive for Rab11a or Rab35 at a given time point, and the proportion of Rab11a- and Rab35-positive CCVs in a population of infected cells vary over time. previous studies including ours have only observed stable fusion events between the CCV and host endosomes and lysosomes [14, 20, 21, 58]. The dynamic nature of CCV-Rab11a/Rab35 interactions could be a result of two possibilities: first, only a subset of CCVs preferentially interacts with the recycling endosomes at a given time point and the number of preferred CCVs interacting with a particular Rab protein varies depending on the stage of infection, and second, it is possible that the recycling endosomes make transient contact with the CCVs in a ‘kiss-and-run’ manner [59], to release their content into the CCV and/or to receive recyclable cargo from the CCV and then dissociate. However, further experimentation is required to test these hypotheses and could be an intriguing area of future investigation.

Our observation that *Coxiella* infection led to an increased recycling endosome content at 3 dpi builds upon the previous observation that *Coxiella* expands the endosomal compartment in infected cells [56]. It is possible that increased number of recycling endosomes accumulate more Tf488 in *Coxiella*-infected cells observed in that study [56]. Interestingly, in our study, we did not observe an increase in the recycling endosome content at 6 dpi. This may be a result of an already low number of endosomes in the *Coxiella-*infected cells at 6 dpi [21]. Nonetheless, these data, taken together with our previous data indicate that *Coxiella* has a dual effect on host endosomes: while it reduces the number of mature endosomes in infected cells [21], *Coxiella* simultaneously elevates the number of recycling endosomes during infection. While the full characterization of these regulations requires further testing, we propose a model that *Coxiella* reroutes the ‘immature’ endosomes from the degradative pathway into the recycling pathway, potentially protecting the host cell from the detriments of the ‘immature’ endosomes.

Our siRNA-mediated knockdown experiments revealed that Rab11a and Rab35 are essential for CCV expansion at 3 and 6 dpi, respectively. A recent study with Rab11a knockout HeLa cells revealed that Rab11a deficiency increases the number of late endosomes and lysosomes within the cells [60]. It is possible that in our experiments, Rab11a knockdown led to an increased number of lysosomes that is detrimental to CCV acidity and *Coxiella* growth. Future research analyzing the lysosomal pH and enzyme activity in *Coxiella-*infected, Rab11a/Rab35 knockdown cells will help further characterize the CCV-recycling endosome interactions.

Overall, our data revealed novel host-*Coxiella* interactions that appear to be beneficial for CCV maturation. Future research into characterizing the role of Rab11a and Rab35 in *Coxiella* pathogenesis, and the molecular mechanisms of *Coxiella* enhancement of recycling endosome content will potentially identify novel therapeutic targets for *Coxiella*.

## 5 Acknowledgments

We thank Stacey Gilk (University of Nebraska Medical Center) for providing the parental HeLa cells and mCherry-*Coxiella burnetii*. We are also thankful to Mary Weber (University of Iowa) for providing the EGFP-tagged Rab plasmids. We thank Manan Damani (Midwestern University) for assistance with confocal and fluorescent microscopy. We also thanks Vijitha Kantety and Ruthvik Gundala for helpful suggestions.

